# Longitudinal neuromelanin changes in prodromal and early Parkinson’s disease in humans and rat model

**DOI:** 10.1101/2024.10.22.619619

**Authors:** Jean-Baptiste Pérot, Anthony Ruze, Rahul Gaurav, Sana Rebbah, Capucine Cadin, Arnaud Le Troter, Lucas Soustelle, Laura Mouton, Romain Valabrègue, Annabelle Parent, Graziella Mangone, François-Xavier Lejeune, Isabelle Arnulf, Jean-Christophe Corvol, Marie Vidailhet, Mathieu D. Santin, Miquel Vila, Stéphane Lehéricy

## Abstract

Studies in animal models of Parkinson’s disease (PD) suggested that the accumulation of the neuromelanin (NM), a pigment contained in nigral dopaminergic neurons, could trigger neurodegeneration above a pathogenic threshold. Here we investigated this hypothesis using NM-sensitive MRI in rodents and in patients with isolated rapid eye movement sleep disorders (iRBD) subjects, a prodromal phase of parkinsonism, and early PD. We first combined NM-sensitive MRI and histology to study NM accumulation and neurodegeneration in a humanized rat model of PD. NM-MRI signal changes were biphasic with an initial increase due to the accumulation of NM in dopaminergic neurons, followed signal decrease due to neurodegeneration. In healthy subjects and patients with iRBD, NM-MRI signal increased initially and then decreased similarly as in rodents after reaching a similar maximum signal intensity in both groups. In early PD and converted iRBD patients, NM-MRI signal drop was greater than in healthy individuals. Results in animals and humans show that NM-sensitive MRI is a marker of the intracellular NM accumulation up to a threshold then of neuronal degeneration beyond this threshold and agree with the hypothesis of a pathogenic threshold of NM triggering neurodegeneration.

## Introduction

Preferential degeneration of dopaminergic neurons in the substantia nigra (SN) and reduction in striatal dopamine are key features of Parkinson’s disease (PD)^1^. Dopaminergic neurons of the SN, and other PD-vulnerable catecholaminergic neurons such as noradrenergic neurons of the locus coeruleus (LC), contain a polymeric pigment called neuromelanin (NM). Although a pathological hallmark of PD is the accumulation of misfolded, aggregated α-synuclein, which is a major component of Lewy bodies, in the brain and in particular the SN^1^, several lines of evidence suggest that NM may also play an important role in PD pathogenesis and in the preferential vulnerability of catecholaminergic neurons. In the SN of PD patients, neuronal loss occurs in melanized neurons and cell loss directly correlates with the percentage of NM-pigmented neurons^2^. Studies have suggested a dual role of NM, protective at the early stage by binding toxins and redox active metals^3^, and toxic once these systems have been exhausted^4^. In addition, neuroinflammatory changes in PD are highly localized within NM-containing areas^1^ and extracellular NM released from dying neurons can activate microglia^5,6^. NM is also associated with α-synuclein that redistributes to the lipid component of NM at early PD stages^7^ and becomes entrapped within NM granules^8^. However, because NM is absent in most animal species, including rodents, its potential role in PD pathogenesis has not been experimentally studied in vivo until the recent development of NM-producing animal models overexpressing human melanin-producing enzyme tyrosinase (hTyr) via an adeno-associated virus (AAV) vector^9,10^. AAV-hTyr-injected animals presented progressive intracellular build-up of human-like NM in the SN, with intracellular concentration reaching similar levels in 2 months old rats^9,11^ and 4 months old macaques^10^ than in PD patients, ultimately compromising neuronal function and viability. These results raised the hypothesis of a pathogenic threshold above which intracellular NM accumulation triggers in an age-dependent manner the main pathological features of PD, including motor deficits, Lewy pathology and nigrostriatal neurodegeneration^12^. This hypothesis was further supported by histological analyses of the SN of subjects with PD and incidental Lewy body disease who had increased intracellular NM levels in dopaminergic neurons compared with age-matched healthy controls^9^.

The most commonly used neuroimaging technique able to detect neuromelanized structures in the human brain is NM-sensitive MRI (NM-MRI). The origin of the NM-MRI signal is still debated but probably includes the T_1_ reduction effect of NM, which is paramagnetic when bound to iron to form NM-iron complexes^13,14^, and the high density of water protons in the nucleus^15,16^. NM-MRI has been proposed as a potential biomarker of melanized neurons^17–19^ and hence of nigral degeneration^14,20,21^. Histological studies in the human SN found that NM appeared at around 2-3 years, accumulated with age, and then, depending on the study, remained stable after the second decade of life^22^ or increased subsequently^23^. This evolutionary profile was clarified in healthy subjects using NM-MRI which showed a progressive increase in the NM signal until reaching a plateau around 50-60 years of age followed by a decrease beyond that age^24^, in line with the concept of a pathogenic threshold of NM accumulation. The authors hypothesized that combined effects of intracellular NM accumulation^25^ and age-related neuronal death could explain this behavior, although this remains to be proven. NM-MRI contrast-to-noise ratio (CNR) is reduced in the SN and LC of PD patients and patients with isolated rapid eye movement (REM) sleep behavior disorder (iRBD), a prodromal condition of parkinsonism^26–28^. Previous studies have suggested that these changes occur after a long prodromal phase preceding motor symptom onset by 10 to 20 years for putaminal dopamine uptake^29,30^ and by 5 to 6 years for SN volume loss assessed using NM-MRI^27,31^. In PD, death of the dopaminergic neurons in the SN is accelerated^32^, and recent work suggested that NM accumulation could also be accelerated^9^.

The hypothesis of a pathogenic threshold of NM accumulation, possibly even in healthy aging individuals, can thus be studied using NM-MRI techniques. Characterizing the NM-MRI contrast changes versus age and disease duration in normal and pathologic conditions could thus enable to better understand the pathogenic mechanisms triggering neurodegeneration in nigral dopaminergic neurons. Furthermore, if the characterization of the evolution profile and the pathological threshold of NM before the onset of the disease is not possible to date in PD, this study is possible in the AAV-hTyr rat model and in patients at prodromal stages of parkinsonism, e.g. iRBD. Here, we used NM-MRI in the AAV-hTyr rat model and in human healthy and PD subjects to answer the following questions: 1) In the AAV-hTyr rat model of PD, what is the temporal evolution of the NM-MRI signal and 2) can we explain the modifications of this NM signal over time by the successive combination of NM accumulation followed by neurodegeneration of melanized neurons beyond a pathological NM threshold in vivo? 3) In healthy human subjects, do we reproduce such an evolution of the NM signal with a maximum around 50-60 years and a decline beyond this age? 4) In patients with iRBD, is the NM threshold the same as in controls and how does NM evolve before and after conversion to parkinsonism? 5) In patients with iRBDs after conversion, is the slope of NM-MRI signal loss similar to that of PD patients? In the present work, we used longitudinal quantitative MRI to characterize the NM-MRI contrast in vivo in the AAV-hTyr rat model, with histological confirmation, and we then compared these results with longitudinal NM-MRI from clinical dataset including healthy volunteers (HV), iRBD, and PD patients.

## Materials and methods

### Animal study

#### Animals

Forty 1-month-old CD male rats were received from Charles Rivers Laboratories. After 2 weeks of acclimatation, rats were handled daily for 2 weeks to reduce the stress due to experimentation. Animals were housed by pairs with 12h day/night cycle, water and food ad libitum, and nesting enrichment that was changed once a week, together with litter.

We conducted a longitudinal imaging study with this cohort of rats. MRI protocol was realized 2 weeks prior to the injection of AAV-hTyr as a baseline, and 1, 2, 4, and 8 months following the injection. Animals were 2 months old at the time of injection. 10 rats were euthanized after each time point for histological evaluation. Rats were randomly dispatched for euthanasia between timepoints before the start of the study. **Figure 1** summarizes the study design.

**Figure 1:**
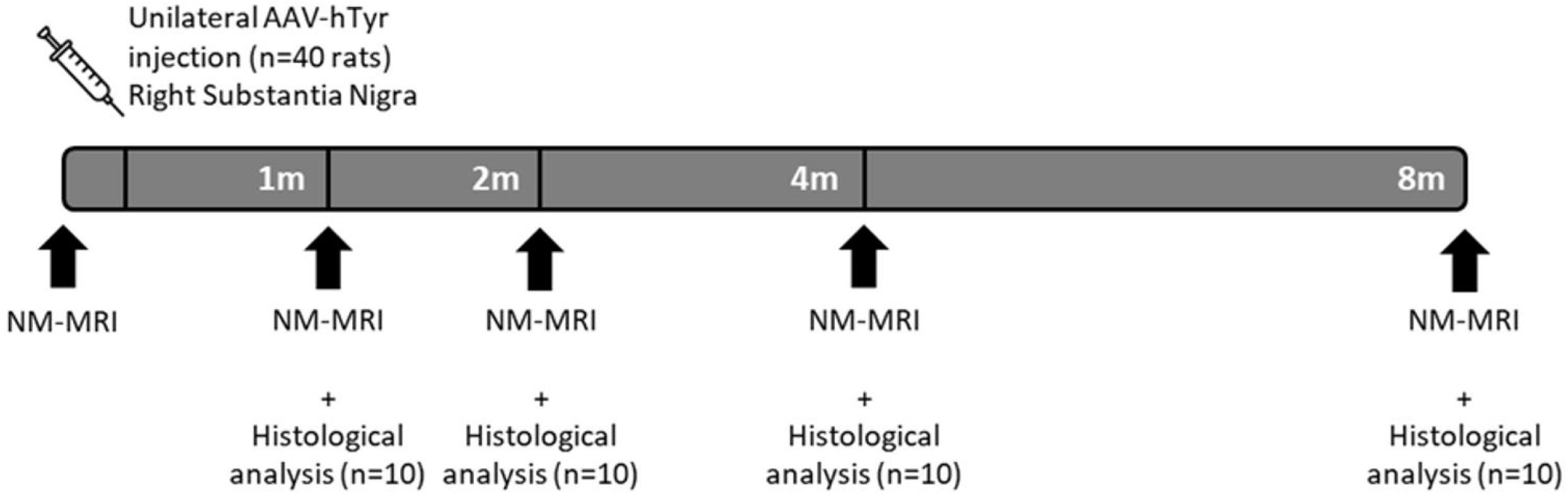
Study design of the longitudinal imaging experiment on AAV-hTyr rats cohort.

#### Surgical procedures

Recombinant AAV vector serotype 2/1 expressing the human tyrosinase cDNA driven by the cytomegalovirus (CMV) promoter (AAV-hTyr; concentration 9,48E+12 gc/mL) were produced at the Viral Vector Production Unit of the Autonomous University of Barcelona (UPV-UAB, Spain). After the first MRI experiment, all rats were injected with AAV-hTyr in the right SN only. Detailed procedures are available in Supplementary Information.

#### Preclinical MRI protocol

Rats were anesthetized with inhalation of isoflurane (3% for induction, 1.5-2% for maintain) and placed inside a 11.7T magnet (Bruker BioSpec; Bruker BioSpin, Ettlingen, Germany) with a birdcage resonator and a receive only head surface coil. The body temperature was monitored and regulated at 37 degrees Celsius during the whole procedure, including induction of anesthesia. A 2D axial T_1_-weighted NM-MRI sequence was acquired after installation as well as three 3D multi gradient echo sequences with varying flip angles (FA) and MT preparation for quantitative MRI (T_1,_ T_2_*, QSM, MPF). A transmit field (B_1_^+^) map was acquired for bias correction using the Actual Flip Imaging method^33,34^. Power calibration was performed before each sequence to account for heating of electronic components. 2D multi-echo spin-echo sequence was added to the protocol when possible for T_2_ mapping. Parameters of acquisitions are detailed in Supplementary Information.

#### Image processing

Rat brain images were reconstructed from raw data using in-house MATLAB program. T_1_ mapping was computed from MTOFF_FA6 and MTOFF_FA24 images using the Variable Flip Angle method^35^. R_2_* was estimated by exponential fitting of the signal from the 16 echoes of MTOFF_FA6 image. QSM and MPF were computed from the three 3D multi gradient echo sequences (Supplementary Information).

Multi-contrast template was then computed from MTOFF_FA6, R_1_, R_2_*, MPF, and QSM baseline images from 24 animals using antsMultivariateTemplateConstruction.sh from ANTS^36^. Following construction of the global template, a subject-specific template was computed for each subject using images from the different timepoints.

The ipsilateral SN was manually delineated at 1mpi on individual NM-MRI images in native space. This segmentation was co-registered to the global template and flipped on the left-right axis for segmentation of the contralateral SN pars compacta (SNpc). Ipsi- and contralateral segmentations were then coregistered to subject-specific template and to images of all time points using ANTS.

#### Tissue collection

10 animals were euthanized after each MRI session for brain tissue collection. Rats were euthanized and perfused with 4% paraformaldehyde before brain extraction. Brains were then post-fixed for 3 days, cryoprotected, frozen and stored at −80°C until slicing with cryostat.

#### Immunostaining

Immunostaining of TH was realized on brain slices using the MULTIVIEW PLUS IHC Kit (Enzo Life Science). Quantification of intracellular neuromelanin was realized with slides mounted in Mowiol 4-88 aqueous solution directly after unfreezing and 3 rinses in distilled water, with no staining.

#### Cell counting and extracellular NM quantification

After drying of mounting medium, slides were scanned using an Olympus Slideview VS200 slide scanner together with the Olyvia 3.3 software to obtain 20x high resolution micrographs of the samples. These were then analyzed by a specific artificial intelligence (AI)-assisted algorithm implemented as in ^37^ for the quantification of TH^+^/NM^+^cells and extracellular NM aggregates.

For all quantifications, SNpc was manually drawn in both hemispheres on 12 TH-immunostained brain slices covering the whole region. As some brain slices may have been damaged and discarded, total counts were normalized by the number of retained slices. Animals were entirely discarded from the analysis when there was less than 7 retained slices. The ratio of quantified cells in ipsilateral compared with contralateral SNpc was calculated for each animal. For TH+ cells, we divided the number of cells ipsilateral by the number of cells contralateral to have a percentage of cells ipsilateral compared to contralateral.

### Human study

#### Participants

All subjects were prospectively recruited in the longitudinal, prospective, observational, case-control ICEBERG study (ClinicalTrials.gov Identifier: NCT02305147, ethics committee approval: IRB of Paris VI, RCB 2014-A00725-42) at the Paris Brain Institute. This cohort included healthy volunteers (HVs), iRBD patients and PD patients assessed using MRI at baseline (V1) and after 2 years (V2) and 4 years (V4). Patient inclusion criteria comprised a clinical diagnosis of iRBD or PD, respectively performed by sleep neurologists and movement disorder specialists, 18–75 years of age, minimal or no cognitive disturbances (Mini-Mental State Examination score > 26/30) and, for PD patients, time elapsed from the first appearance of motor symptoms (i.e., disease duration) < 4 years. Clinical examination is detailed in Supplementary Information. Patients with PD met the UK Parkinson’s disease Society Brain Bank criteria^38^. Patients with iRBD had a history of dream-enacting behaviours with (potentially) injurious movements and enhanced tonic chin muscle tone or complex behaviours during REM sleep but did not meet the criteria for PD or dementia^39^. All participants gave written informed consent. The demographic and clinical characteristics of the subjects are presented in Table 1.

**Table 1.**
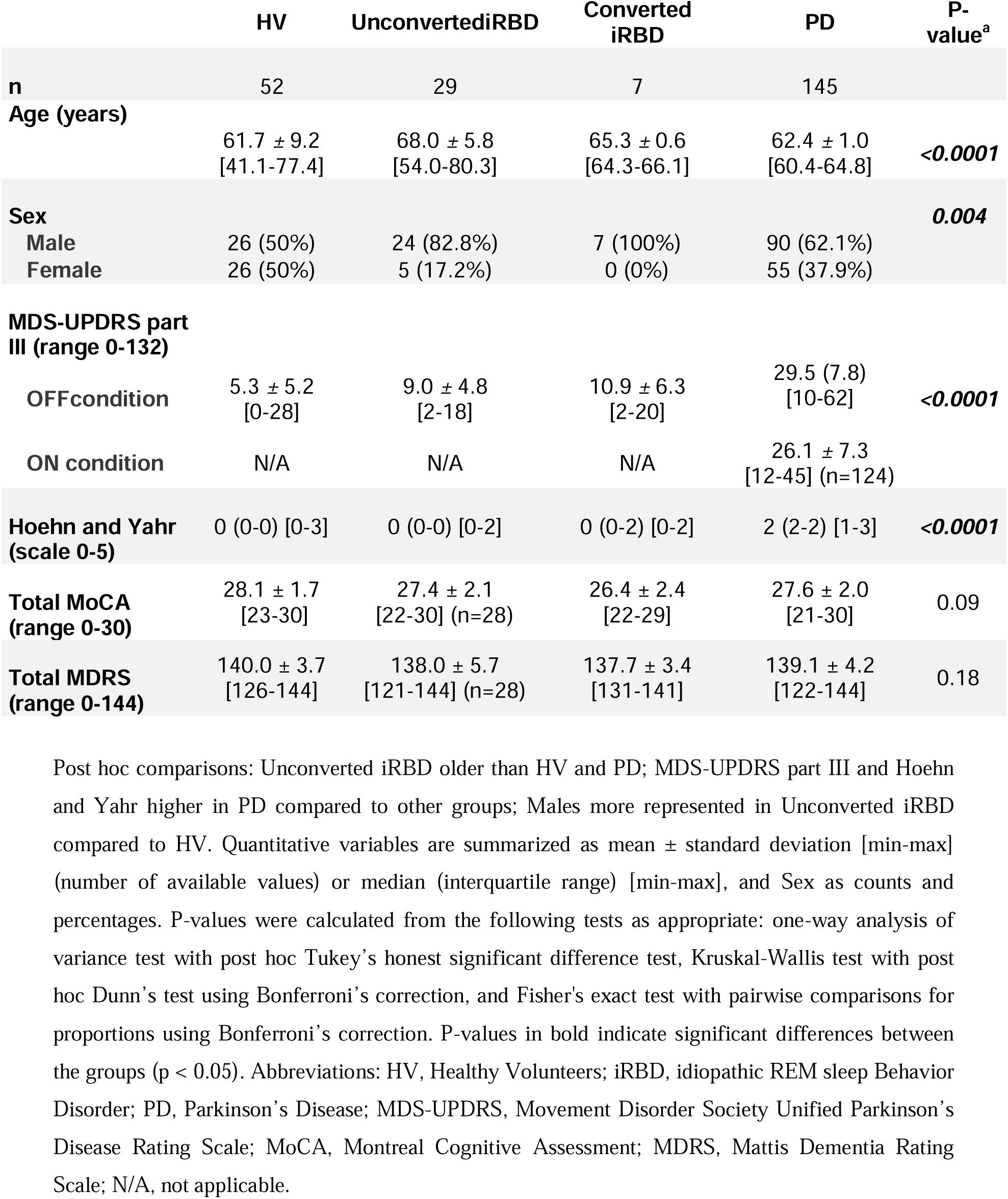
Demographic and clinical characteristics at the baseline visit (V1).

#### Clinical MRI protocol

MRI was performed for all subjects on a Siemens Prisma 3T scanner (Siemens Healthineers) with a 64-channel head reception coil. Whole brain anatomic image was acquired with T_1_-weighted 3D MP2RAGE. NM-sensitive MRI was acquired using T_1_-weighted 2D turbo spin echo as previously described^30^.

Image co-registrations were performed using NiftyReg (v1.5.58). Region of interest segmentations were delineated using the FreeSurfer image viewer (v5.3.0).

Brain extraction and segmentation into grey matter, white matter and CSF was performed on whole-brain anatomical images from MP2RAGE sequence using Statistical Parametric Mapping software for MATLAB (SPM12).

NM-sensitive turbo spin echo images of the SN were coregistered to anatomical images using affine transformation. The SN was then manually segmented by two trained raters blind to the groups, as previously described^27,40^. SN volume was extracted and corrected by total intracranial volume. Normalized Signal Intensity (NSI) of the SN was computed in comparison with a background region including the tegmentum and superior cerebral peduncle.

### Statistical analyses

All statistical analyses were conducted using R version 4.2.2 (R Development Core Team, 2022). R_1_, R_2_, R_2_*, MTR, MPF were included in the statistical analysis as the change rate between ipsilateral and contralateral as following (S_ipsi_-S_contra_)/S_contra_. CNR was calculated from NM-MRI image as (S_ipsi_-S_contra_)/σ_background_. QSM was analyzed as (S_ipsi_-S_contra_) without normalization due to values very close to zero. Intracellular and extracellular NM were included as raw data, neurodegeneration as n_ipsi_/n_contra_ ratio.

Longitudinal data were analyzed by fitting separate Linear Mixed Models (LMMs) to each imaging modality. To compare histological measurements over time, which consisted in independent observation points between the time points, we performed one-way analysis of variance (ANOVA) with time from injection as a factor. Correlations between the changes in NM-MRI and histological quantification were examined using Spearman’s rank correlation coefficients (Spearman’s rho).

The level of statistical was set at p or adjusted p < 0.05 (two-sided) for all tests.

### Study approval

All animal studies were conducted according to the French regulation (EU Directive 2010/63/EU – French Act Rural Code R 214-87 to 126). The animal facility was approved by veterinarian inspectors (authorization B-751319) and complies with Standards for Humane Care and Use of Laboratory Animals of the Office of Laboratory Animal Welfare (CEEA - 005). All procedures received approval from the ethical committee (APAFIS # 34975-2022012412181911 v9).

## Results

### AAV-hTyr rats

#### Age-dependent NM accumulation and neurodegeneration in AAV-hTyr rats

Previous studies showed that overexpression of hTyr in rat and non-human primate SN resulted in age-dependent production of human-like NM within nigral dopaminergic neurons associated with nigrostriatal neurodegeneration, Lewy body-like pathology and a PD phenotype^9–11^. In this rat model, animals showed a steady increase in intracellular NM levels in dopaminergic neurons over time associated with neurodegenerative changes that started by 1 month post-AAV injection (mpi) but became significant by 4mpi^9^. Here, we first sought to reproduce these findings and then to characterize by NM-MRI the longitudinal changes in NM accumulation and neurodegeneration in the SN of AAV-hTyr rats.

One-month-old male rats received a single unilateral stereotaxic injection of AAV-hTyr above the right SN as described previously^9^. As expected, following injection, rats showed a progressive accumulation of intracellular (**Figure 2A and 2D**) and extracellular (**Figure 2B and 2E**) NM in the ipsilateral SN. At 1mpi intracellular NM optical density (OD) was already significantly different from zero, considered as the theoretical baseline level as regular rats lack NM (*p*<0.01, one-sample, one-sided t-test). Intracellular NM OD was further increased at 4mpi compared with 1mpi (+48.7%, *p*<0.05) and at 8mpi compared with other time points (+73.9% vs 1mpi, *p*<0.001). In parallel to NM accumulation, nigral dopaminergic neurons started to degenerate in AAV-hTyr rats, as previously reported^9^. At 1mpi, AAV-hTyr rats already showed a non-significant tendency to dopaminergic neuronal loss, with a 21.7% reduction in the ratio of the number of tyrosine hydroxylase-positive (TH+) neurons in the injected (i.e. melanized) vs contralateral (i.e. non-melanized) hemisphere (**Figure 2C and 2F**). The number of TH+ cells was significantly reduced in the SN in the hemisphere ipsilateral to AAV-hTyr injection as shown by the decrease in the ratio of the number of TH+ neurons in the ipsilateral hemisphere divided by the number of TH+ neurons in the contralateral hemisphere at 4mpi (−68.4%, *p*<0.001) and 8mpi (−74.8%, *p*<0.001) (**Figure 2C**). Following degeneration, NM from dying melanized neurons is released to the extracellular space, both in humans and AAV-hTyr rats^9^. In agreement with this, extracellular aggregates appear more numerous at 2mpi than at 1mpi, although there is no significant main effect of time for extracellular NM (+40.0%, *p*=0.3) (**Figure 2B and 2E**).

**Figure 2:**
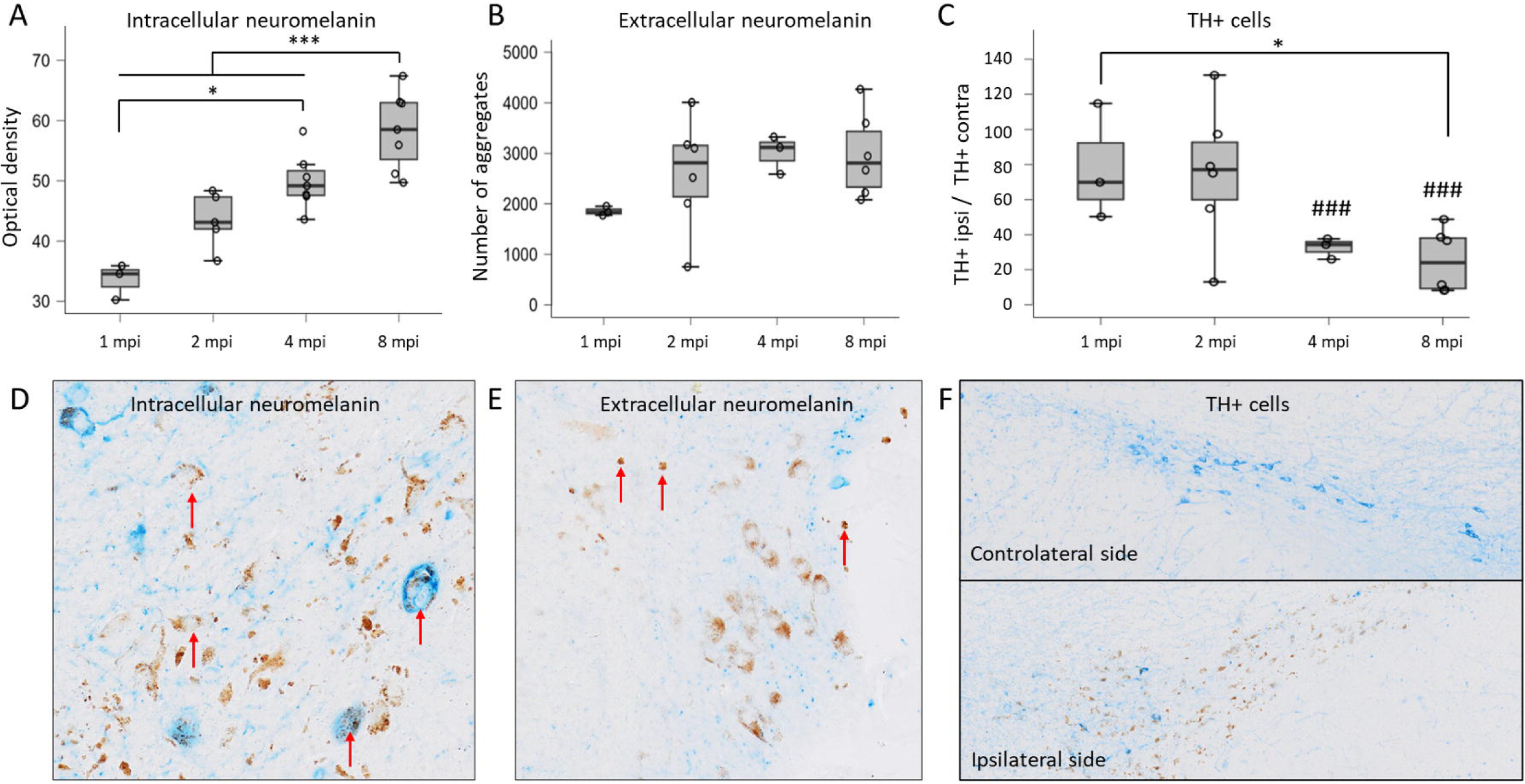
NM accumulation and neurodegeneration following AAV-hTyr injection in rat SN. **A)** Optical density of intracellular NM in ipsilateral SN of rats euthanized at different months post injection (mpi) (1mpi: 4 rats, 2mpi: 7 rats, 4mpi: 8 rats, 8mpi: 7 rats). **B)** Number of extracellular NM aggregates in ipsilateral SN of rats euthanized at different mpi (1mpi: 3 rats, 2mpi: 6 rats, 4mpi: 3 rats, 8mpi: 5 rats). **C)** Ratio of the number of TH+ cells in the ipsilateral (TH+ipsi) compared to the contralateral SN (TH+contra) in rats euthanized at different mpi (1mpi: 3 rats, 2mpi: 6 rats, 4mpi: 3 rats, 8mpi: 6 rats). \**p*<0.05, \*\*\**p*<0.001 (between time points), ###*p*<0.001 (ipsilateral vs contralateral). One-way ANOVA with Tukey’s post-hoc test. Histological images of the SN ipsilateral to AAV-hTyr injection with TH staining showing **D)** intracellular NM (arrows) and **E)** extracellular NM (arrows). **F)** Histological images of the SN contralateral and ipsilateral to AAV-hTyr injection with TH staining showing reduced TH+ neurons (blue) in the injected ipsilateral side and NM (brown color).

#### Biphasic age-dependent NM-MRI signal changes in AAV-hTyr rats

We next characterized the temporal evolution of the NM-MRI signal in AAV-hTyr rats. As histological analyses showed an accumulation of intracellular NM followed by a reduction of NM-containing neurons, we would expect NM signal intensity to follow a biphasic curve, with an initial signal increase followed by a secondary decrease after reaching a maximum threshold.

AAV-hTyr rats were scanned longitudinally at baseline, 1, 2, 4 and 8mpi using an 11.7T MRI system (**Figure 3A**). As expected, AAV-hTyr rats showed an increase in ipsilateral SN NM-MRI contrast-to-noise ratio (CNR) between baseline and 1mpi (x11.58, *p*<0.001) followed by a progressive decrease in CNR from 2 to 8mpi (**Figure 3B**). CNR was still higher at 2mpi (x8.26, *p*<0.001) and 4mpi (x4.58, *p*<0.05) compared to baseline. At 8mpi, CNR was no longer significantly different from baseline levels. Compared to 1mpi, CNR was significantly decreased at 2mpi (−26.4%, *p*<0.05), 4mpi (−55.7%, *p*<0.001) and 8mpi (−63.0%, *p*<0.001). CNR was also significantly decreased at 4mpi (−39.7%, *p*<0.05) and 8mpi (−49.7%, *p*<0.05) compared with 2mpi. These results are in line with the concept that degeneration of melanized neurons occurs beyond a pathological NM threshold in vivo.

**Figure 3:**
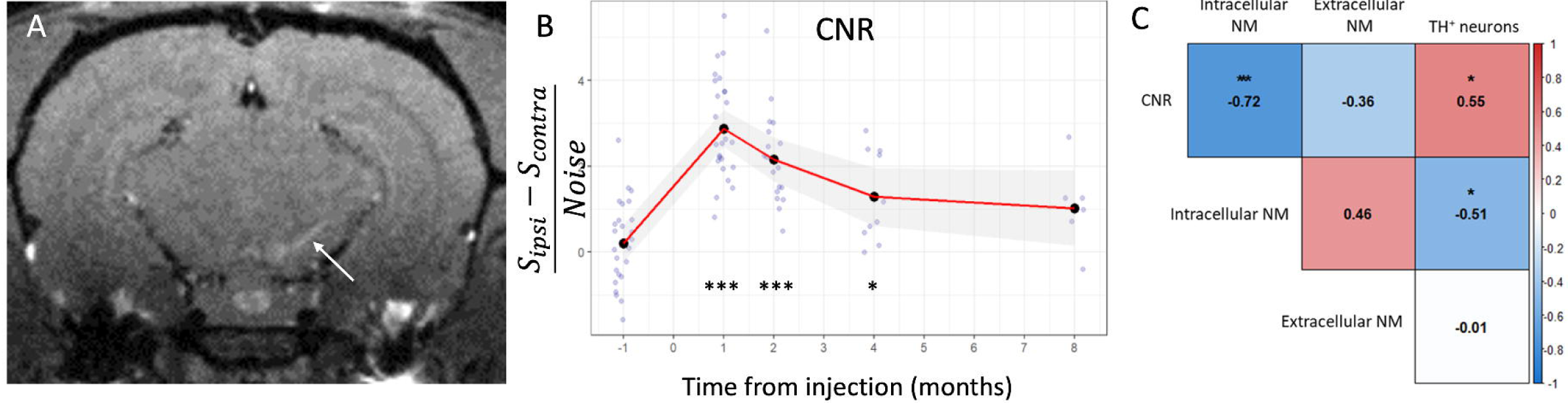
Progression of NM contrast and correlation with NM accumulation and neurodegeneration. **A)** Representative NM-MRI image of an AAV-hTyr-injected rat at 1 mpi. Melanized SN is detected as an ipsilateral hyperintense area (arrow). **B)** Contrast-to-Noise Ratio between the ipsilateral and contralateral SN as function of time from AAV-hTyr injection. Black dots represent mean value per timepoint. Red line and grey shading represent estimated marginal means and confidence intervals. \**p*<0.05 and \*\*\**p*<0.001 versus baseline, Linear Mixed Model followed by post-hoc pairwise comparisons using the FDR correction. S_ipsi_ = signal in the ipsilateral SN, S_contra_ = signal in the contralateral SN. **C)** Correlation matrix of CNR parameter with histological quantification between 1 mpi and 8 mpi Spearman’s rho correlation coefficient. Significant correlations are highlighted. \**p*<0.05, \*\*\**p*<0.001.

Overall, our results showed that age-dependent NM-MRI signal changes in AAV-hTyr rats were first associated with NM accumulation and then with neurodegeneration of NM-containing neurons. The initial increase of NM-MRI CNR (**Figure 3B**) paralleled the accumulation of intracellular (**Figure 2A**) and extracellular NM (**Figure 2B**) in these animals. From 1mpi onwards, decreasing NM-MRI CNR signal positively correlated with the decreasing number of nigral dopaminergic neurons (*r*=0.55, *p*<0.05), **Figure 3C**). In turn, the number of dopaminergic neurons correlated negatively with intracellular NM (r=-0.51, p<0.05). Indirectly, this led to negative correlation between NM density and CNR during this degenerative phase (**Figure 3C)**. These results suggest that in absence of overt neurodegeneration (approximately during the first two mpi), signal intensity was mainly driven by the increase in intracellular NM levels. Beyond this stage, after reaching a pathogenic NM threshold, decreasing CNR signal was mainly driven by the loss of NM-containing neurons. Of note, the contribution of extracellular NM to the overall NM-MRI signal seems to be negligible, as extracellular NM levels are maintained stable histologically from 2mpi onwards while NM-MRI signal progressively decreases in parallel to cell loss independently of extracellular NM accumulation.

#### NM-MRI is associated with T_1_ reduction in AAV-hTyr rats

Although NM-MRI signal hyperintensity in humans is clearly observed in regions that contain melanized catecholaminergic neurons (notably SN and LC), the exact origin of the observed signal is still debated. Two hypotheses are most often put forward. The most widespread hypothesis attributes the elevation of the T_1_ signal to the paramagnetic properties of NM when it is associated with metals such as iron^14^. The main competing hypothesis is that the main source of NM-MRI contrast is the high water content of catecholaminergic neurons^15,16^. Notably several studies have reported that magnetization transfer (MT) preparation increased the contrast between the SN and the surrounding brain tissue, which led several authors to consider that MT was the main source of contrast^41,42^. Our previous results showing an elevation of the T_1_ signal associated with the presence of NM suggested that the elevation of signal was well linked to the presence of NM^30^. To further study the origin of the so-called NM-MRI signal, we studied in AAV-hTyr rats the correlations of this signal with other MRI parameters such as the longitudinal relaxation rate R_1_ (=1/T_1_), the proportion of protons linked to macromolecules, evaluated by the MT ratio (MTR) and the macromolecular proton fraction (MPF, evaluated by the quantitative MT), and the presence of iron in the SN (evaluated by the transverse relaxation rates R_2_ and R_2_*, and quantitative susceptibility mapping QSM) (**Figure 4A-C**).

**Figure 4:**
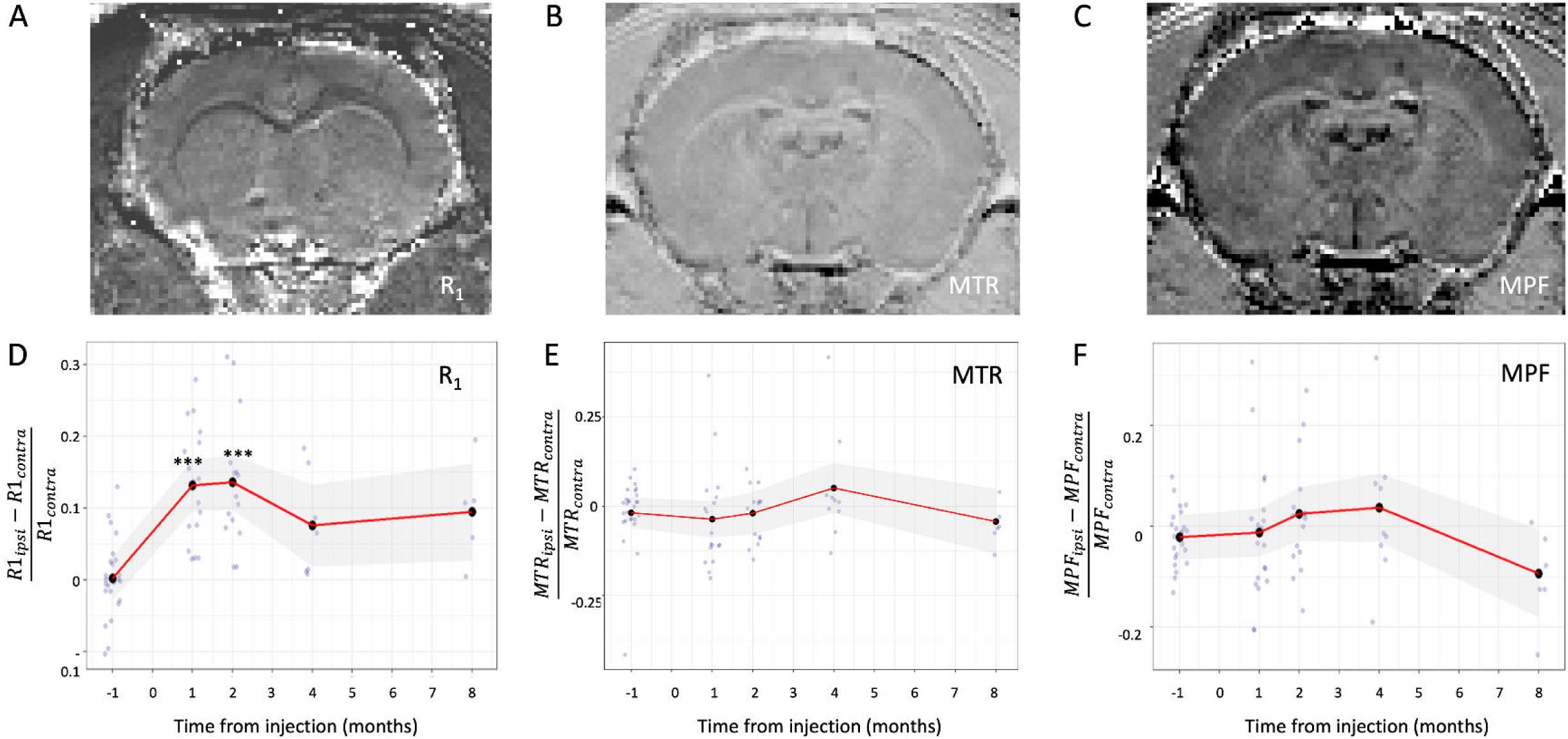
Progression of magnetization transfer and longitudinal relaxation parameters in time. **A-C)** Representative brain images of **A)** R_1_, **B)** MTR and **C)** MPF. **D-F)** Variation over time from AAV-hTyr injection of **D)** R_1_, **E)** MTR, and **F)** MPF parameters in the ipsilateral SN compared with the contralateral SN. Black dots represent mean value per time point. Red line and grey shading represent estimated marginal means with 95% confidence intervals. \*\*\**p*<0.001 versus baseline, LMM followed by post-hoc pairwise comparisons using the FDR correction.

The variation over time of quantitative R_1_ mapping between ipsilateral and contralateral SN was similar to that of NM-MRI CNR (**Figure 4D**), highlighting the contribution of T_1_ reduction effect of NM to the NM-MRI contrast. R_1_ in the ipsilateral SN was significantly higher at 1mpi (+13.2%, *p*<0.001) and 2mpi (+14.7%, *p*<0.001) compared with the contralateral SN. In contrast, neither MTR nor MPF were significantly sensitive to NM accumulation (**Figure 4E-F**). NM-MRI CNR correlated significantly with R_1_ (*r*=0.63, *p*<0.001) but not with MPF or MTR. MTR was strongly correlated with MPF (*r*=0.85, *p*<0.001) and R_1_ (r=-0.7, *p*<0.001).

#### Iron-sensitive magnetic susceptibility changes in AAV-hTyr rats

Iron changes were assessed using QSM, R_2_* and R_2_ mapping (**Figure 5A-C**). Magnetic susceptibility (QSM) increased progressively in AAV-hTyr rats (**Figure 5D**), suggesting iron accumulation in the ipsilateral SN of these animals. Magnetic susceptibility difference between ipsilateral and contralateral SN was significantly increased between the baseline and 1mpi (x3.28, *p*<0.05), and 2, 4, and 8mpi (x5.94, x7.26, and x7.87 respectively, *p*<0.001). Variation of R_2_* (**Figure 5E**) and R_2_ over time (**Figure 5F**) between the ipsilateral and contralateral SN were not significantly different. Magnetic susceptibility was also correlated positively with R_2_ (*r*=0.66, *p*<0.05).

**Figure 5:**
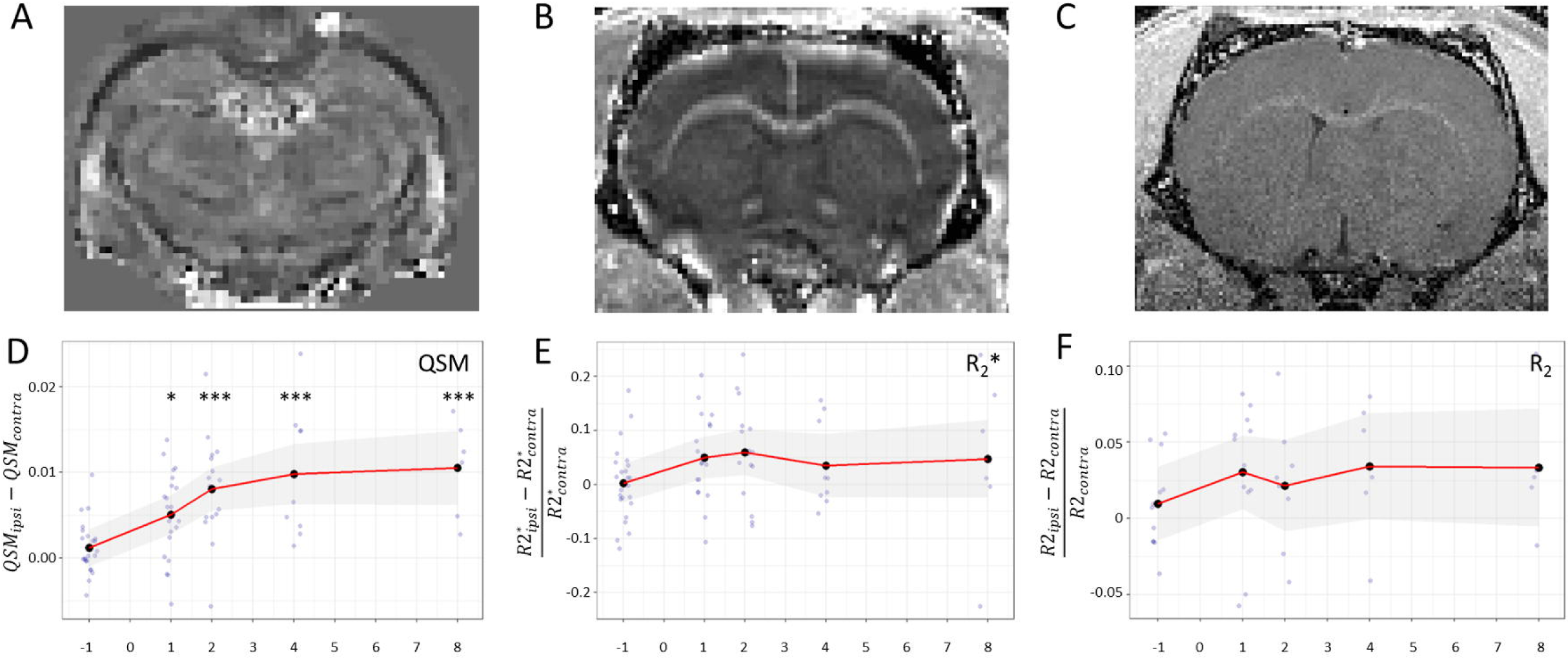
Progression of susceptibility and transverse relaxation parameters in time. **A-C)** Representative brain images of **A)** QSM, **B)** R_2_*and **C)** R_2_. **D-F)** Correlation of magnetic susceptibility parameters with CNR in AAV-hTyr rats. **D)** Differences in QSM between ipsilateral and contralateral SN over time from injection. Variations in **E)** R_2_* and **F)** R_2_ parameters in the ipsilateral compared with the contralateral SN over time from injection. Black dots represent mean value per timepoint. Red line and grey shading represent estimated marginal means with 95% confidence intervals. \**p*<0.05 and \*\*\**p*<0.001 versus baseline, LMM followed by post hoc pairwise comparisons using the FDR correction.

In summary, our results suggest that NM-MRI signal changes observed in the SN of AAV-hTyr rats are related to the changes in T_1_ relaxation time induced by the production of NM and not to changes in the macromolecular proton fraction of the tissue. In addition, these changes are associated with an increase in iron concentration in the SN of these animals.

### Longitudinal NM-MRI in human subjects

#### Age-dependent NM-MRI signal changes in healthy volunteers and patients with iRBD

The results in the humanized NM-producing rat model demonstrated that the MRI signal sensitive to NM was indeed a marker of the presence of NM in the SN of these animals up to a certain threshold, and then reflected the decreasing number of melanized dopaminergic neurons beyond this threshold. In addition, NM-MRI signal in this model was directly linked to the presence of NM and not to the concentration of protons bound to macromolecules. These results may help us to better interpret the NM-MRI signal variations observed in humans. In healthy subjects, NM-MRI studies have shown an increase in the signal with age reaching a plateau around 50-60 years of age^24^ corresponding to an accumulation of NM in histological analyses^25^, then a decrease beyond that age. We therefore first studied the variations of the NM-MRI signal in the SN of 52 healthy volunteers (HV) recruited in the ICEBERG study at the Paris Brain Institute. Subjects were aged between 41 and 77 years and had no current or prior history of psychiatric or neurological disorders. Longitudinal follow-up of the HV over a period of up to 4.5 years showed a parabolic inverted u-shape trajectory of the NM-MRI signal intensity in the SN with a maximum signal intensity of 100.2 reached at 54.9 years of age, followed by a decrease in signal intensity beyond this threshold (**Figure 6**, black line) in line with a previous study^24^.

**Figure 6:**
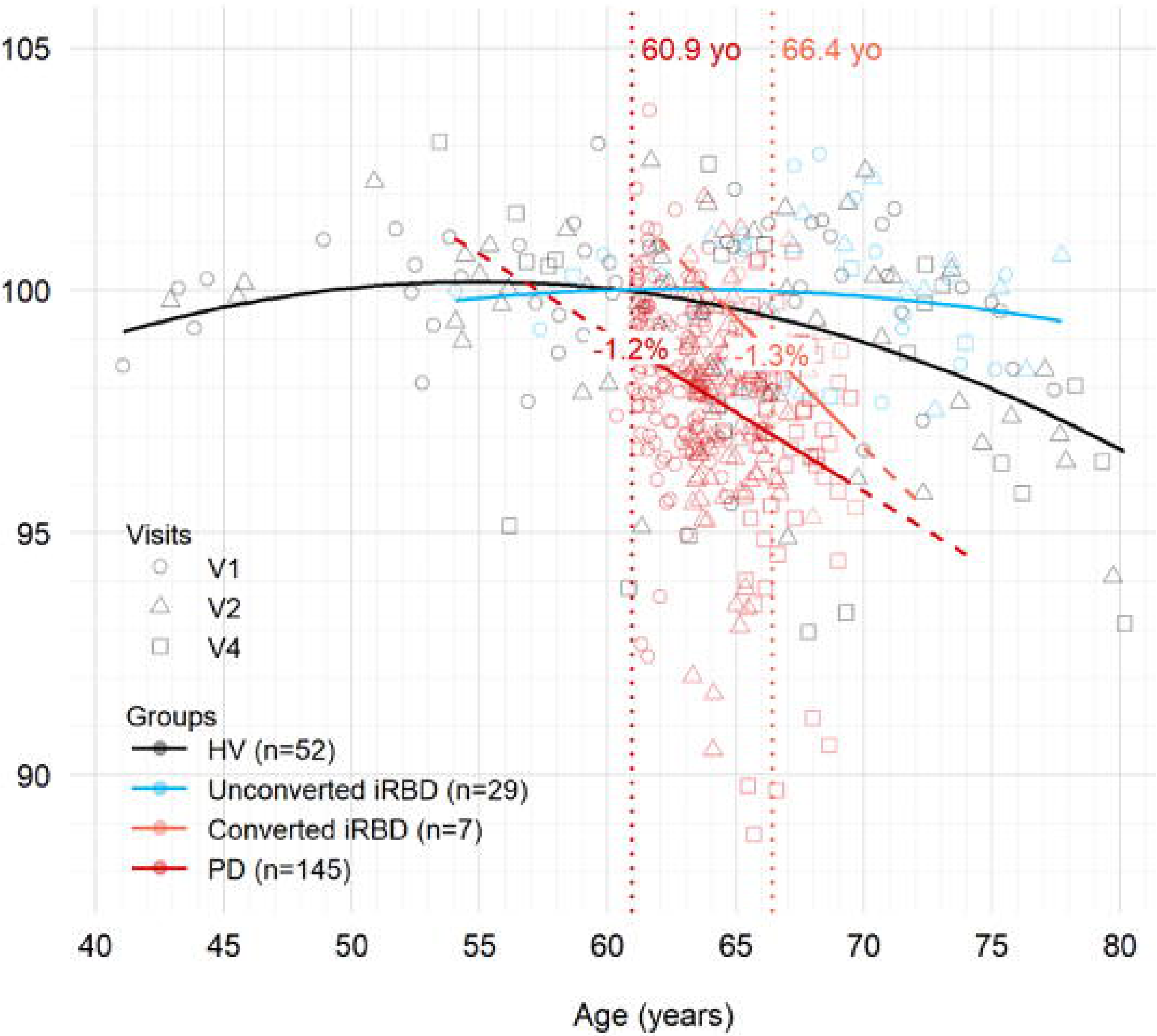
NM-MRI normalized signal intensity (NSI) in the SN as a function of age in years. In HV (black), unconverted iRBD patients (cyan), converted iRBD patients (orange) and PD patients (red). NSI values were obtained by dividing each value by the mean NSI value (110.9) of the control subjects at the baseline visit (V1). Vertical dotted lines represent the mean ages of onset in PD (red) and converted iRBD (orange). For the sake of comparison between groups and within the PD group, all PD patients’ ages were aligned at baseline with the mean age of onset to study the effect of disease duration as the actual age was meaningless. Solid curves were added to indicate the NSI evolution of each group using the “geom_smooth” function of the ggplot2 R package with either a linear fit for PD and converted iRBD patients or a quadratic fit for HV and unconverted iRBD patients. Dashed lines represent extrapolated prolongation of linear trajectory outside of observed age intervals. NSI loss in percentage (%) as indicated by the regression lines at the mean ages of onset was indicated for PD and iRBD converted patients. Abbreviations: NSI, Normalized Signal Intensity; yo, years old; HV, Healthy Volunteers; iRBD, idiopathic REM sleep Behavior Disorder; PD, Parkinson’s Disease.

We then wanted to compare this evolution profile of the NM-MRI signal to that of patients at the prodromal stage of parkinsonism, i.e. iRBD, to determine whether this threshold was similar or increased compared to HV and at what age it was reached. We studied the signal evolution profile of 36 patients with polysomnography-confirmed iRBD included in the ICEBERG study and aged from 54 to 80 years. We found that iRBD patients who did not progress to parkinsonism during the study (n=29 non converted iRBD patients) showed a parabolic trajectory similar to that of HV, delayed by 7.9 years, with a maximum of 100.0 reached at 62.8 years (**Figure 6**, blue line). The threshold appeared therefore the same as in HV.

#### Age-dependent NM-MRI signal changes in converted iRBD and PD patients

Lastly, we analysed the NM-MRI evolution profile in 7 iRBD patients who converted to parkinsonism during the study and compared this profile to that of 145 idiopathic PD patients and HV from the ICEBERG study. In converted iRBD patients (**Figure 6**, orange line) the mean age of conversion was 66.4 years. At the time of conversion, iRBD patients had a 1.3% reduction in NM-MRI signal intensity compared to unconverted iRBD patients. Longitudinal progression of the NM-MRI signal was best fitted by a linear estimation (*r*=-0.537, *SE*=0.056, *p*=0.10). The back projection of this estimation met the curve of unconverted iRBD at the maximum (62.8 years).

The mean age at diagnosis in PD patients was 60.9 years, 5.5 years earlier than that of converted iRBD patients. At the time of onset, PD patients had a 1.2% reduction of NM-MRI signal intensity compared to the mean signal intensity in HV (**Figure 6**, red line) similar to that of converted iRBD patients. Longitudinal imaging of PD patients showed a decrease in NM-MRI signal intensity after onset. This decrease was better fitted by a linear estimate (*r*=-0.326, *SE*=0.326, *p*<0.0001), accelerated compared to HV whose decrease rate is about 0.0007 for the year following the PD onset. The back projection of the linear estimation met the HV curve near maximum (56.8 years). There was no significant difference between the CNR decrease in PD and converted iRBD (*r*=-0.211, *SE*=0.331, *p*=0.53).

Overall, the results in humans support the hypothesis that beyond a certain threshold of NM in the SN, neurodegeneration occurs in HV, iRBD and PD patients and that this threshold appears similar in both healthy and diseased groups. However, neurodegeneration progresses faster in patients and is delayed by approximately 5.5 years in converting iRBD compared with idiopathic PD patients.

## Discussion

This is the first longitudinal study tracking NM-MRI signal changes from the presymptomatic phase to the PD stage with histological comparison in a humanized rat model producing NM and in prodromal parkinsonism. The results showed that NM-sensitive imaging made it possible to follow the progression of NM accumulation in the SN with a first initial phase of progressive increase in the levels of NM in nigral neurons until reaching a specific threshold followed by neurodegeneration beyond this threshold. The results obtained in humans show a similar signal evolution profile in healthy subjects confirming previous report^24^ but also in parkinsonian patients at the prodromal stage of iRBD, although delayed by 7.9 years. In both the healthy and unconverted iRBD groups, the NM-MRI signal increased to a similar peak intensity in both groups, likely reflecting NM accumulation up to a similar intracellular content in dopamine neurons. Beyond this threshold, the signal intensity began to decrease, probably reflecting the occurrence of neurodegeneration. The NM-MRI signal drop linked to neurodegeneration was more accentuated in iRBD patients who had converted to PD and in patients with PD than in healthy individuals.

### Biological origin of the NM-MRI contrast in the humanized rat model of PD

Many studies have already reported a relationship between NM-MRI based signal intensity and the presence of NM in the human SN^14,17–20,43^. In the AAV-hTyr rat and macaque models, the accumulation of NM in the SN was associated with an increase in NM-MRI signal intensity^9,10^. In human specimens, previous studies showed that the area of high signal intensity in NM-MRI images matched with the location of the SN pars compacta^18,44,45^. Positive correlation of NM-MRI signal intensity was reported with the density of NM-containing neurons in 12 regions of interest from the SN of 3 subjects^17^ and with the regional NM concentration from midbrain sections of 7 individuals with PD or PD-related syndromes^19^. The results of the present study complement these data by precisely quantifying the longitudinal changes in NM-MRI signal in relation to NM accumulation in the SN in histological sections. NM-MRI CNR in the ipsilateral SN of AAV-hTyr rats showed a large and rapid increase after injection of the viral vector. At this stage, CNR increase was concomitant with the accumulation of intracellular NM without overt neuronal loss. Following this initial increase, CNR reached a plateau between 1 and 2mpi during which the intracellular NM density continued to increase, the population of TH+ neurons started to decrease by approximately 20-25%, and the accumulation of extracellular NM debris released from dying neurons was established. During this plateau phase, the increase in NM levels and cell loss compensated each other, explaining the relative stability of the signal. From 2mpi onwards, CNR decreased continuously in parallel to cell death, reaching 80% reduction compared to the contralateral SN at 8mpi. During this phase, the decrease in NM-MRI CNR despite the continuous increase in intracellular NM indicates that the reduction in the number of NM-containing neurons at regional level predominated over the progressive increase of NM at cellular level within the remaining, yet-to-degenerate neurons. Together, these results demonstrate that NM-MRI can follow these two successive phenomena.

### Longitudinal progression of NM-MRI signal in healthy aged individuals

Together with previous histological data in humans, our results in the rat model of NM production allowed us to better interpret the evolution of the NM-MRI signal in healthy subjects. On the one hand, our results in HV confirmed those of a previous study^24^ by showing an increase in the NM-MRI signal up to a maximum plateau between 50 and 60 years followed by a signal drop beyond that age. On the other hand, results confirmed the hypothesis that the initial NM-MRI signal increase in HV was due to the age-dependent increase in intracellular NM levels and that the following NM-MRI signal decrease was secondary to the appearance of overt age-related neurodegeneration. Our results are in agreement with histological studies in aging humans, which showed that NM increased continuously through life^23^ and that age-related loss of pigmented neurons occurred even in the absence of clinical PD^46,47^. Our results thus support the hypothesis that loss of pigmented neurons occurs when a maximum threshold of intracellular NM is reached^12^. In healthy aging individuals, this threshold is reached around fifty, according to NM-MRI, an age at which increased NM levels and neurodegeneration compete to explain the MRI signal of NM.

### Longitudinal progression of NM-MRI signal in PD

In idiopathic PD patients, interpolations of NM-MRI signal changes suggested that the pathogenic NM threshold was reached at the same age as in HV but that neurodegeneration was much more marked with an accelerated signal loss slope. In PD patients, additional genetic and/or environmental factors, in combination with pathogenic NM levels, might thus intervene to explain the faster degeneration of vulnerable melanized neurons once the threshold is reached. In PD, the temporal profile of nigrostriatal neurodegeneration has been most often described by a negative mono-exponential function by histological studies^48,49^, studies using radiotracers targeting the dopaminergic system^29,41,50–54^ or by NM-MRI^27,30^. Some studies however have reported a linear decrease in dopamine function with advancing disease duration using DAT-SPECT^55–57^ or NM-MRI^31^.

### Longitudinal progression of NM-MRI signal in iRBD

In unconverted iRBD subjects, the CNR described a similar inverted U-shape curve to that of HV, reaching the same maximum threshold but delayed by 7.9 years. Moreover, iRBD patients who converted to PD during the study showed accelerated decrease in NM-MRI signal like PD patients but delayed by the same number of years. The later onset of NM-MRI signal changes in iRBD was consistent with the known later onset of parkinsonism in iRBD compared to idiopathic PD^58^. This delay suggests a different mechanism of progression of SN neurodegeneration between PD and iRBD patients. The majority of iRBD patients convert to parkinsonism within 10–15 years, including not only PD but also dementia with Lewy bodies and rarely multiple system atrophy, with a phenoconversion rate of approximately 6–8% per year^38,58^. There is strong evidence that iRBD also represents distinct, more severe subtypes of alpha-synucleinopathies^59^. In addition, the pattern of progression of the neurodegenerative process differs between iRBD and PD patients, being more variable in PD patients in which the first neurodegenerative changes occur either in the brainstem or the olfactory areas^60–62^. In the present study, the slope of the NM-MRI signal decrease seemed greater in iRBDs than PD, but the difference was not significant, which could be due to the low number of converted iRBD subjects.

### Pathogenic threshold of NM accumulation in the SN

Together, our results are in line with the hypothesis of an intracellular NM pathogenic threshold contributing to triggering neurodegeneration^12^, both linked to aging and in PD, being accelerated in the latter case. In rodents, hTyr overexpression produced NM, whose accumulation above a specific threshold compromised neuronal function and triggered neurodegeneration replicating a previous study^9^. Interestingly, NM intracellular concentration correlated negatively with neuronal loss in these animals suggesting a relationship between these two phenomena. Supporting this concept, reduction of intracellular NM accumulation in AAV-hTyr rats, either by boosting NM cytosolic clearance with autophagy activator TFEB or by delaying age-dependent NM production through VMAT2-mediated enhancement of dopamine vesicular encapsulation, resulted in a major attenuation of their neurodegenerative phenotype, both at the behavioral and neuropathological level^9,37^. In humans, the average NM signal intensity in PD and converted iRBD patients crossed the parabolic curves of HV and non-converting iRBD at their maximum, respectively, a maximum that was similar between both groups. Together, these results are in line with the hypothesis of a pathogenic threshold of NM concentration and of a role of NM-induced toxicity in PD pathophysiology^12^.

### Physical origin of the NM-MRI contrast

The origin of the NM-MRI contrast is debated. It is usually attributed to T_1_ shortening of NM when bound to metals and particularly iron^13,14^. Such effects may be mostly attributable to iron bound to NM, forming paramagnetic complexes, at least *ex vivo*^13^. Alternatively, recent work has suggested that the high signal intensity observed in the locus coeruleus containing melanized noradrenergic neurons was unlikely due to the presence of unique T_1_-shortening molecules such as NM, but could instead be attributed to the high proton density of water whose T_1_ was shortened by paramagnetic ions^16^. Low MPF due to poor myelination of the SN may also be involved^63^, in which case NM-MRI hyperintensity would not actually come from NM but from a reduced MPF^15,43^. MT experiments also point towards a suppressing effect of signal from the surrounding tissues^13^. In this context, our results indicate that the increase in NM-MRI signal in AAV-hTyr rats was mainly related to the accumulation of intracellular NM but not to MPF or MTR changes, which showed no significant temporal variations. Thus, in this model, NM-MRI signal changes were clearly associated with NM deposition in the SN and the results did not support a strong contribution of macromolecules exchanging with water or the regional proton density of the structures.

### Iron accumulation in the SN of AAV-hTyr rats

Iron-sensitive QSM increased over time in the SN of AAV-hTyr rats. QSM and R_2_* increases have both been associated with iron accumulation with QSM being more sensitive than R_2_* to detect iron changes^64,65^. In particular, QSM has been reported to be sensitive to NM-iron complexes^66^, which represent the main storage system of iron molecules in the SN pars compacta, potentially protecting dopaminergic neurons from iron toxicity^67,68^. Conversely, R_2_ correlates with ferric iron deposits ^66^ and was not significantly changed in AAV-hTyr rats.

### Study limitations

This study presents several limitations. First, rat and human MRI data were not acquired at the same magnetic field, with strong differences in relaxometry parameters. Thus, while based on the same sequence structure, the origin of the NM-MRI contrast at 11.7T may not entirely be the same as 3T. Second, the brain structure and metabolism between the two species are not identical and the phenotype induced by unilateral nigral AAV-hTyr injections in rats do not replicate all the complexity of PD. However, the similarities between PD and the PD-like changes reported previously in AAV-hTyr rats^9^, combined with the changes in the NM-MRI profiles observed here, suggested that results obtained in AAV-hTyr rats could indeed serve to better understand changes observed in humans. Third, the low number of iRBD patients who converted to parkinsonism may have decreased the accuracy of modeling NM-MRI signal loss in this group. However, the results obtained were in agreement with literature data, which reported a later onset of motor disorders in iRBD patients compared to PD patients^58^. Fourth, the relatively short follow-up of the patients did not make it possible to precisely determine the type of decrease in the NM-MRI signal, i.e. mono-exponential or linear. However, this was not the focus of our study and a linear vs exponential fit would not significantly change the results reported here. Fifth, the SN was manually segmented in both rats and humans. Automatic segmentation could allow better reproducibility and avoid potential biases. Lastly, it would have been interesting to include an additional earlier time point (i.e. 0.5mpi) in our AAV-hTyr rat studies, to observe the increase of CNR before neurodegeneration started. However, in practice it was not possible to scan so many animals in such a short time.

### Conclusion

Longitudinal, quantitative *in vivo* characterization of the NM-MRI contrast in the AAV-hTyr rat model allowed the identification of the biological and physical origins of this signal. NM-MRI contrast was associated with T_1_ reduction and was sensitive to NM accumulation in the first phase, and then to neurodegeneration of pigmented dopaminergic neurons. These results supported the hypothesis of a pathogenic threshold of intracellular NM accumulation, followed with iron accumulation. This concept was also consistent with the temporal slope of NM-MRI in the SN of PD patients and may explain the late onset of PD conversion in iRBD patients. More studies are needed to better understand the different progression patterns in iRBD and PD.

## Data availability

Data is available to download in supporting data.

## Supporting information

ARRIVE Author Checklist

STROBE Checklist

## Acknowledgements

The authors would like to thank the HISTOMICS and PHENOPARC platforms of the Paris Brain Institute for their support. The B_1_^+^ mapping SPGR prototype sequence used in experiments was provided by the CRMBM laboratory (Aix Marseille Univ, CNRS, CRMBM, Marseille, France).

## Supplementary material

Supporting Data

Supplementary Information

