## Supplementary material for "Longitudinal neuromelanin changes in prodromal and early Parkinson’s disease in humans and rat model": ARRIVE Author Checklist

NOTE: Please save this file locally before filling in the table, DO NOT work on the file within your internet browser as changes will not be saved. Adobe Acrobat Reader (available free here) is recommended for completion.

### The ARRIVE guidelines 2.0: author checklist

#### The ARRIVE Essential 10

These items are the basic minimum to include in a manuscript. Without this information, readers and reviewers cannot assess the reliability of the findings.

| Item | | Recommendation | Section/line number, or reason for not reporting |
|---|---|---|---|
| **Study design** | 1 | For each experiment, provide brief details of study design including: | |
| | | a. The groups being compared, including control groups. If no control group has been used, the rationale should be stated. | Figure 1 |
| | | b. The experimental unit (e.g. a single animal, litter, or cage of animals). | Figure 1 |
| **Sample size** | 2 | a. Specify the exact number of experimental units allocated to each group, and the total number in each experiment. Also indicate the total number of animals used. | Figure 1 |
| | | b. Explain how the sample size was decided. Provide details of any *a priori* sample size calculation, if done. | Figure 1 |
| **Inclusion and exclusion criteria** | 3 | a. Describe any criteria used for including and excluding animals (or experimental units) during the experiment, and data points during the analysis. Specify if these criteria were established *a priori*. If no criteria were set, state this explicitly. | M&M - Cell counting and extracellular NM quantification |
| | | b. For each experimental group, report any animals, experimental units or data points not included in the analysis and explain why. If there were no exclusions, state so. | Supporting Data |
| | | c. For each analysis, report the exact value of *n* in each experimental group. | Figures legends |
| **Randomisation** | 4 | a. State whether randomisation was used to allocate experimental units to control and treatment groups. If done, provide the method used to generate the randomisation sequence. | M&M - Animals |
| | | b. Describe the strategy used to minimise potential confounders such as the order of treatments and measurements, or animal/cage location. If confounders were not controlled, state this explicitly. | N.A. |
| **Blinding** | 5 | Describe who was aware of the group allocation at the different stages of the experiment (during the allocation, the conduct of the experiment, the outcome assessment, and the data analysis). | Supplementary Information - Intracellular NM quantification |
| **Outcome measures** | 6 | a. Clearly define all outcome measures assessed (e.g. cell death, molecular markers, or behavioural changes). | All figures, supporting Data, M&M - Statistical Analyses |
| | | b. For hypothesis-testing studies, specify the primary outcome measure, i.e. the outcome measure that was used to determine the sample size. | M&M - Statistical Analyses |
| **Statistical methods** | 7 | a. Provide details of the statistical methods used for each analysis, including software used. | M&M - Statistical Analyses, Supplementary Information - Statistical Analyses |
| | | b. Describe any methods used to assess whether the data met the assumptions of the statistical approach, and what was done if the assumptions were not met. | M&M - Statistical Analyses, Supplementary Information - Statistical Analyses |
| **Experimental animals** | 8 | a. Provide species-appropriate details of the animals used, including species, strain and substrain, sex, age or developmental stage, and, if relevant, weight. | M&M - Animals |
| | | b. Provide further relevant information on the provenance of animals, health/immune status, genetic modification status, genotype, and any previous procedures. | M&M - Animals |
| **Experimental procedures** | 9 | For each experimental group, including controls, describe the procedures in enough detail to allow others to replicate them, including: | Supplementary Information - Surgical Procedure |
| | | a. What was done, how it was done and what was used. | |
| | | b. When and how often. | |
| | | c. Where (including detail of any acclimatisation periods). | |
| | | d. Why (provide rationale for procedures). | |
| **Results** | 10 | For each experiment conducted, including independent replications, report: | |
| | | a. Summary/descriptive statistics for each experimental group, with a measure of variability where applicable (e.g. mean and SD, or median and range). | All figures |
| | | b. If applicable, the effect size with a confidence interval. | Figures legends |
