## Supplementary material for "Longitudinal neuromelanin changes in prodromal and early Parkinson’s disease in humans and rat model": STROBE Checklist

**STROBE statement: Reporting guidelines checklist for cohort, case-control and cross-sectional studies**

| **SECTION** | **ITEM NUMBER** | **CHECKLIST ITEM** | **REPORTED ON PAGE NUMBER:** |
| --- | --- | --- | --- |
| **TITLE AND ABSTRACT** |  |  |  |
|  | 1a | Indicate the study’s design with a commonly used term in the title or the abstract | 1 |
|  | 1b | Provide in the abstract an informative and balanced summary of what was done and what was found | 1 |
| **INTRODUCTION** |  |  |  |
| Background and objectives | 2 | Explain the scientific background and rationale for the investigation being reported | 2-4 |
|  | 3 | State specific objectives, including any pre-specified hypotheses | 4 |
| **METHODS** |  |  |  |
| Study design | 4 | Present key elements of study design early in the paper | 4,7 |
| Setting | 5 | Describe the setting, locations, and relevant dates, including periods of recruitment, exposure, follow-up, and data collection | 7 |
| Participants | 6a | Cohort study—Give the eligibility criteria, and the sources and methods of selection of participants. Describe methods of follow-up  Case-control study—Give the eligibility criteria, and the sources and methods of case ascertainment and control selection. Give the rationale for the choice of cases and controls  Cross-sectional study—Give the eligibility criteria, and the sources and methods of selection of participants | 7 |
|  | 6b | Cohort study—For matched studies, give matching criteria and number of exposed and unexposed  Case-control study—For matched studies, give matching criteria and the number of controls per case  Variables |  |
| Variables | 7 | Clearly define all outcomes, exposures, predictors, potential confounders, and effect modifiers. Give diagnostic criteria, if applicable | 7-8 |
| Data sources/measurements | 8* | For each variable of interest, give sources of data and details of methods of assessment (measurement). Describe comparability of assessment methods if there is more than one group. | Supporting Data |
| Bias | 9 | Describe any efforts to address potential sources of bias. | 8 |
| Study size | 10 | Explain how the study size was arrived at | 7 |
| Quantitative variables | 11 | Explain how quantitative variables were handled in the analyses. If applicable, describe which groupings were chosen and why . | 8 |
| Statistical methods | 12a | Describe all statistical methods, including those used to control for confounding | 8 |
|  | 12b | Describe any methods used to examine subgroups and interactions | 8 |
|  | 12c | Explain how missing data were addressed | Supplementary Information |
|  | 12d | Cohort study—If applicable, explain how loss to follow-up was addressed  Case-control study—If applicable, explain how matching of cases and controls was addressed  Cross-sectional study—If applicable, describe analytical methods taking account of sampling strategy | 7 |
|  | 12e | Describe any sensitivity analyses |  |
| **RESULTS** |  |  |  |
| Participants | 13a | Report numbers of individuals at each stage of study—eg numbers potentially eligible, examined for eligibility, confirmed eligible, included in the study, completing follow-up, and analysed | Figures legends |
|  | 13b | Give reasons for non-participation at each stage |  |
|  | 13c | Consider use of a flow diagram |  |
| Descriptive Data | 14a | Give characteristics of study participants (eg demographic, clinical, social) and information on exposures and potential confounders | Table 1 |
|  | 14b | Indicate number of participants with missing data for each variable of interest | Supporting data |
|  | 14c | Cohort study—Summarise follow-up time (eg, average and total amount) | 7 |
| Outcome Data | 15* | Cohort study—Report numbers of outcome events or summary measures over time  Case-control study—Report numbers in each exposure category, or summary measures of exposure  Cross-sectional study—Report numbers of outcome events or summary measures | 7 |
| Main Results | 16a | Give unadjusted estimates and, if applicable, confounder-adjusted estimates and their precision (e.g. 95% confidence interval). Make clear which confounders were adjusted for and why they were included | All figures |
|  | 16b | Report category boundaries when continuous variables were categorized |  |
|  | 16c | If relevant, consider translating estimates of relative risk into absolute risk for a meaningful time period |  |
|  | 16d | Report results of any adjustments for multiple comparisons | 8 |
| Other Analyses | 17a | Report other analyses done—e.g. analyses of subgroups and interactions, and sensitivity analyses |  |
|  | 17b | If numerous genetic exposures (genetic variants) were examined, summarize results from all analyses undertaken |  |
|  | 17c | If detailed results are available elsewhere, state how they can be accessed |  |
| **DISCUSSION** |  |  |  |
| Key Results | 18 | Summarise key results with reference to study objectives | 14 |
| Limitations | 19 | Discuss limitations of the study, taking into account sources of potential bias or imprecision. Discuss both direction and magnitude of any potential bias | 17 |
| Interpretation | 20 | Give a cautious overall interpretation of results considering objectives, limitations, multiplicity of analyses, results from similar studies, and other relevant evidence | 14-17 |
| Generalisability | 21 | Discuss the generalisability (external validity) of the study results  Other information | 14-17 |
| **FUNDING** |  |  |  |
|  | 22 | Give the source of funding and the role of the funders for the present study and, if applicable, for the original study on which the present article is based | 20 |

*Give information separately for cases and controls in case-control studies and, if applicable, for exposed and unexposed groups in cohort and cross-sectional studies.
