## Supplementary Information for "Longitudinal neuromelanin changes in prodromal and early Parkinson’s disease in humans and rat model"

**Surgical procedures**

Anesthesia was induced by inhalation of 3% isoflurane and maintained with 1.5-2% isoflurane. Rats received subcutaneous injection of Buprenorphine at 0.1mg/kg for analgesia 20 min prior to injection. Then skin and subcutaneous tissue were incised in stereotaxic frame, and skull was perforated 5.2 mm posterior and 2 mm right to the bregma. After perforation, 2µL of AAV-hTyr solution was injected 7.6 mm below the dura mater at a speed of 0.4 µL/min. After injection the subcutaneous tissue and the skin were sutured separately with resorbing thread. The surgical procedure lasted 1h30. Body temperature of the rats was monitored and regulated during the whole procedure. A second injection of Buprenorphine was realized 4-6 hours after the injection. Buprenorphine was mixed to the drinking water for 24h.

**Preclinical MRI protocol**

Preclinical MRI protocol included 2D T_1_ weighted image for neuromelanin imaging (FLASH, TR/TE=333/6.8ms, in-plane resolution=150x150µm², 16 slices, slice thickness=0.4mm, acquisition time=12 min) ; three 3D multi gradient echo sequences with varying flip angles (FA) and MT preparation were acquired for quantitative MRI (TR=47ms, TE_1_:ΔTE:TE_L_=1.71:1.91:30.36ms, 16 echoes, spatial resolution=200µm isotropic, whole brain coverage, t_MT_=10ms, B_1,MTe_^peak^=7.8µT, acquisition time=32 min each). The specific parameters for the three sequences were respectively FA=6°, offset frequency (Δ) of 100kHz (MTOFF_FA6); FA=24°, Δ=100kHz (MTOFF_FA24); FA=6°, Δ=5kHz (MTON_FA6) ; and a 2D Multi Spin Echo sequence for T_2_ mapping (TR/TE=4000/7ms, 8 echoes, in-plane resolution=150x150 µm², slice thickness=400 µm, 50 slices, acquisition time=12min48s, n=12 animals at baseline and 1mpi, n=6 at 2mpi and 4mpi, and n=5 at 8mpi).

**Quantitative Susceptibility Mapping**

For QSM generation procedure, raw data were exported, and complex images (magnitude and phase) were then reconstructed with in-house software in Matlab (Mathworks, USA). The signal magnitude and phase from multiple channels of the receive coil were combined using an in-house optimized method. Then, the local field map was calculated by non-linear fitting of the complex gradient echo signal over echo times^66^, background magnetic fields were removed using the Laplacian boundary value method^67^ and the local field-to-magnetic susceptibility inverse problem was solved using the L1-Morphology Enabled Dipole Inversion method^68^.

**Quantitative Magnetization Transfer**

Macromolecular Proton Fraction (MPF) was calculated from MTOFF_FA6, MTON_FA6, T_1_ and B_1_ maps using Yarnykh’s single-point method^69^ with optimized parameters at 11.7T^70^. Python code for MPF estimation can be found at <https://github.com/lsoustelle/qMT>.

**Immunohistochemistry**

Animals were anesthetized with inhalation of isoflurane and injected intraperitoneally with xylazine (500mg/kg) for analgesia. After 15 minutes rats received a lethal dose of Euthasol (500mg/kg) intraperitoneally. After death, perfusion of PBS was performed in the left cardiac ventricle for 6 min, followed with perfusion of 4% paraformaldehyde (PFA). Brain was then extracted and post-fixed for 3 days in 4% PFA. Brains were then washed 3 times with PBS and plunged in a solution of 30% sucrose for cryoprotection. After sinking, brains were frozen in -50°C isopentane and stored at -80°C. Brains were then sliced at -20°C with a cryostat. SN was sampled with 2 series of 10 microscope slides, each containing 6 brain slices 14 µm thick. Slides were stored at -80°C. Slides were dried at room temperature for 30 min and rinsed 2 times in wash buffer. Samples were incubated in Blocker/diluent solution for 10 min and rinsed 2 times in wash buffer. Slides were then incubated overnight at 4°C with primary antibody (1:1000, Calbiochem, Cat#657012) and rinsed 5 min in wash buffer before 30 min incubation with Multiview Plus Rabbit AP secondary antibody solution. After 2 rinses in wash buffer the staining was revealed in anti-rabbit blue AP Chromogen substrate for 10 min at room temperature. Following revelation, samples were rinsed 3 times in wash buffer and 2 times in distilled water and then mounted in Mowiol 4-88 aqueous solution. Quantification of intracellular neuromelanin was realized with slides mounted in Mowiol 4-88 aqueous solution directly after unfreezing and 3 rinses in distilled water, with no staining.

**Intracellular NM quantification**

Intracellular NM density was quantified by measuring optical density of intracellular NM in unstained slices. After drying of mounting medium, slides were scanned using an Olympus Slideview VS200 slide scanner together with the Olyvia 3.3 software to obtain 20x high resolution micrographs of the samples. For each animal, a minimum of 30 cells were sampled in the entire ipsilateral substantia nigra. NM was manually delineated in cells by two different raters blinded to the time of euthanasia. Mean optical density of NM was extracted for each cell and normalized by optical density of the background. Then, mean NM optical density was calculated for each animal by averaging the normalized NM optical density values of all cells.

**Clinical MRI protocol**

MRI was performed for all subjects on a Siemens Prisma 3T scanner (Siemens Healthineers) with a 64-channel head reception coil. Whole brain anatomic image was acquired with T_1_-weighted 3D MP2RAGE (TR/TE=5000/2.98 ms, voxel size=1x1x1 mm^3^, acquisition time=8 min 12 s). NM-sensitive MRI was acquired using T_1_-weighted 2D turbo spin echo (TR/TE=890/13 ms, in-plane resolution=0.3x0.3 mm^2^, 48 slices, slice thickness=3 mm, acquisition time=6 min 55 s).

**Clinical examination**

Clinical examination included the Hoehn and Yahr scale, which evaluates the severity of Parkinson’s disease-related symptoms^68^; the Movement Disorder Society Unified Parkinson’s Disease Rating Scale (MDS-UPDRS) part III in the OFF condition, which evaluates motor disability >12 hours after withdrawing dopaminergic treatment^69^; and the Mattis dementia rating scale (MDRS^70^) and the Montréal cognitive assessment score (MoCA^71^) , which evaluate cognitive impairment.

**Statistical Analyses**

Longitudinal data were analyzed by fitting separate Linear Mixed Models (LMMs) to each imaging modality using the “*lmer*” function in the lme4 package (v1.1-31)^78^. All LMMs included time from injection as independent variable, with a random intercept to account for the repeated measurements per animal. To compare histological measurements over time, which consisted in independent observation points between the time points, we performed one-way analysis of variance (ANOVA) with time from injection as a factor. Significance of all the main effects or the interaction effects were assessed by Type II Wald chisquare tests using the “*Anova*” function in the car package (v3.1-1). Post-hoc pairwise comparisons were then performed on the significant effect based on estimated marginal means from the linear models using the “*emmeans*” function in the emmeans package (V.4.5). The resulting p-values of the pairwise comparisons were adjusted for multiple comparisons using the false discovery rate (FDR) method. Additionally, to examine the effect of laterality, we tested the estimated means of the change rates against zero at each timepoint.

Correlations between the changes in NM-MRI and histological quantification were examined using Spearman’s rank correlation coefficients (Spearman’s rho). Longitudinal association between the MR parameters was analyzed using repeated measures correlations with the rmcorr package (v0.6.0)^79^.

To investigate the age-evolution of the NM signal in the subjects of the ICEBERG cohort, the NM signal data were normalized by dividing all values by the mean value of the HV group calculated at baseline (110.9) and multiplying by 100. For the sake of visualization and comparison, the PD patients’ baseline ages were aligned arbitrarily to the mean age at onset of the PD group. Doing this allowed to consider and compare patients by disease duration rather than their less meaningful actual age. An approximation of the evolution of the NM signal was made graphically on the basis of the cohort follow-up data by fitting the data of each group with a quadratic trend (HV and iRBD) and a linear trend (PD and an isolated fourth group of 7 iRBDs that converted to PD during the follow-up) using the “*geom_smooth*” function of the ggplot2 package (v3.4.2) (Wickham, 2016, [http://ggplot2.org](http://ggplot2.org/)). The degree of decrease in the NM signal for both PD and converted iRBD groups was reported with the slope (r) of the linear regression line and its standard error (SE), and then compared with each other using the “*emtrends*” function in the emmeans package.
